# Evolution and Genetic Characterization of Seoul Virus in Wild Rats *Rattus Norvegicus* from an urban park in Lyon, France 2020-2022

**DOI:** 10.1101/2023.04.12.536564

**Authors:** Hussein Alburkat, Teemu Smura, Marie Bouilloud, Julien Pradel, Gwendoline Anfray, Karine Berthier, Lara Dutra, Anne Loiseau, Thanakorn Niamsap, Viktor Olander, Diana Sepulveda, Vinaya Venkat, Nathalie Charbonnel, Guillaume Castel, Tarja Sironen

## Abstract

Seoul virus (SEOV) is an orthohantavirus primarily carried by rats that can cause hemorrhagic fever with renal syndrome (HFRS) in humans. Nowadays, its incidence is likely underestimated. We developed a comprehensive serological and molecular characterization of SEOV in *Rattus norvegicus* population from a popular urban park within a large city (‘Park of La Tête d’Or’, Lyon, France) between 2020 and 2022. We confirmed the circulation of SEOV in rats from the park (seroprevalence 17.2%). The SEOV strains detected showed high genetic similarity with the strain previously described in 2013 in this area. We found low structuring of wild rat populations within Lyon city. This study confirms the circulation of SEOV in Lyon city, in a park where opportunities for SEOV transmission to humans are numerous. Given the high gene flow between rat populations in the park and the rest of the city, we recommend conducting city-wide SEOV surveillance.

## Introduction

Seoul virus (SEOV) is an orthohantavirus (*Hantaviridae* family) that is primarily carried by rats, particularly the brown rat (*Rattus norvegicus*) and the black rat (*Rattus rattus*).

It is transmitted to humans through contact with infected rats or their droppings, urine, or saliva. In its rodent hosts, SEOV infection is considered persistent and asymptomatic, but in humans it can cause hemorrhagic fever with renal syndrome (HFRS). The severity of the disease can range from mild to severe (1 -2% mortality) (*1,2,3*). Because of the widespread distribution of rats, SEOV circulates worldwide (*4*). It has been detected in Asia, Europe, Africa and the Americas, both in rural and urban regions, although most studies have been conducted in rural regions (*4*). Despite its potentially high pathogenicity, the incidence of SEOV infections remains probably underestimated (*5*). There are several factors that influence the risk of SEOV infection in humans. Among them, urbanization and climate change are critical processes as they have significant impacts on the distribution and expansion of rats worldwide, and on the probability of close contact between rats and humans(*6*) In Europe, SEOV detection (and transmission to humans) has been documented in pet rats (*7*) as well as in wild rat populations (*8, 9, 10*). The strains detected in pet and wild rats are different (*7*) which raises questions about the origins of wild rats’ strains.

The presence of SEOV in France has been detected in both humans and rats previously (*11, 12*). The first SEOV-HFRS case detected in Europe, confirmed by molecular evidence, was found in a pregnant woman from Replonges (*12*), a locality close to Lyon, a densely populated city with over 500,000 inhabitants in France. Since then, several studies have demonstrated SEOV circulation in Lyon city area. Molecular evidence indicated the presence of SEOV in brown rats that were sampled in 2003 from colonies bred specifically for the study of rodenticides in Lyon (*10*). The presence of SEOV was also confirmed by RT-PCR in 18/128 (14%) wild brown rats sampled in 2011 in Lyon (*13*). Complete genome sequences were obtained, which allowed to classify SEOV variants from Lyon within the lineage 7 (*13*).

The persistence and emergence or re-emergence of zoonotic pathogens in urban environments is a major concern for public and veterinary health. Cities and their peripheries are transportation hubs and are undergoing profound and accelerated socio -environmental changes (*14*). This may lead to the profusion of rodents such as rats and mice in cities, and consequently, increased exposure of humans to rodent-borne zoonotic agents such as SEOV.

As only symptomatic treatment is available for hantavirus -associated diseases, preventing infections remains essential. This mainly consists in avoiding contact with rodents, their secretions and excretions or in implementing measures for controlling rodent populations. A better understanding of the ecological, epidemiological and evolutionary processes involved in the transmission of SEOV is required to mitigate the infection risk and to optimize prevention strategies. Further SEOV, genomic sequences are critical for epidemiological surveillance by allowing the tracking of the spread and evolution of the virus. As an orthohantavirus member, the genome structure of SEOV is tri -segmented, with large (L), medium (M), and small (S) segments. These segments encode different viral proteins; L encodes the viral RNA -dependent RNA Polymerase (RdRP), M encodes two envelope glycoproteins (Gn and Gc), and S encodes nucleocapsid protein (N) (*15, 16*). It also has high mutation rates and short generation times (*17, 18*).

We conducted in-depth investigation of SEOV circulation and genetic evolution in Lyon city. We focused on an urban park that is close to the place where SEOV was detected and sequenced from rats trapped in 2010-2012 (*13*). Due to its high popularity, the park is a hotspot for human/animal interactions, which makes the risk of SEOV transmission likely. We characterized SEOV seroprevalence and sequenced complete coding regions in brown rats (*Rattus norvegicus*) surveyed in this park. We compared the SEOV genomes with the genetic data collected in the same area ten years ago to analyse the evolutionary history of SEOV strains in this urban park (*13*). As such, this study provides a comprehensive serological and molecular characterization of SEOV in *Rattus norvegicus* from Lyon. It gives essential information to design surveillance strategies regarding SEOV risk in a popular urban park within a large city.

## Materials and Methods

### Sampling

Rats were collected during spring 2020, spring and autumn 2021 and spring 2022 in the urban park of “la Tête d’Or” in Lyon city (Figure 1). The capture of animals was carried out in various types of habitats (wooded areas, buildings, zoological park, hay storage yards, riverside vegetation, restaurants, kids’ playgrounds, botanical garden and landfills), using live traps baited with sunflower seeds, carrots and sardine. Traps were set in places where animals had been previously detected (Figure 1), and they were removed after 10 or 11 nights. Each trap was geolocated. The traps were checked daily early in the morning. Animal dissections and measurements were performed according to the protocols described in (*19*). Capture data were registered in the small mammal database (BPM, http://bpm-cbgp.science), and the associated biological samples (such as organs, blood and parasites) were included in the CBGP reference collection of small mammals (*20*).

**Figure 1.**
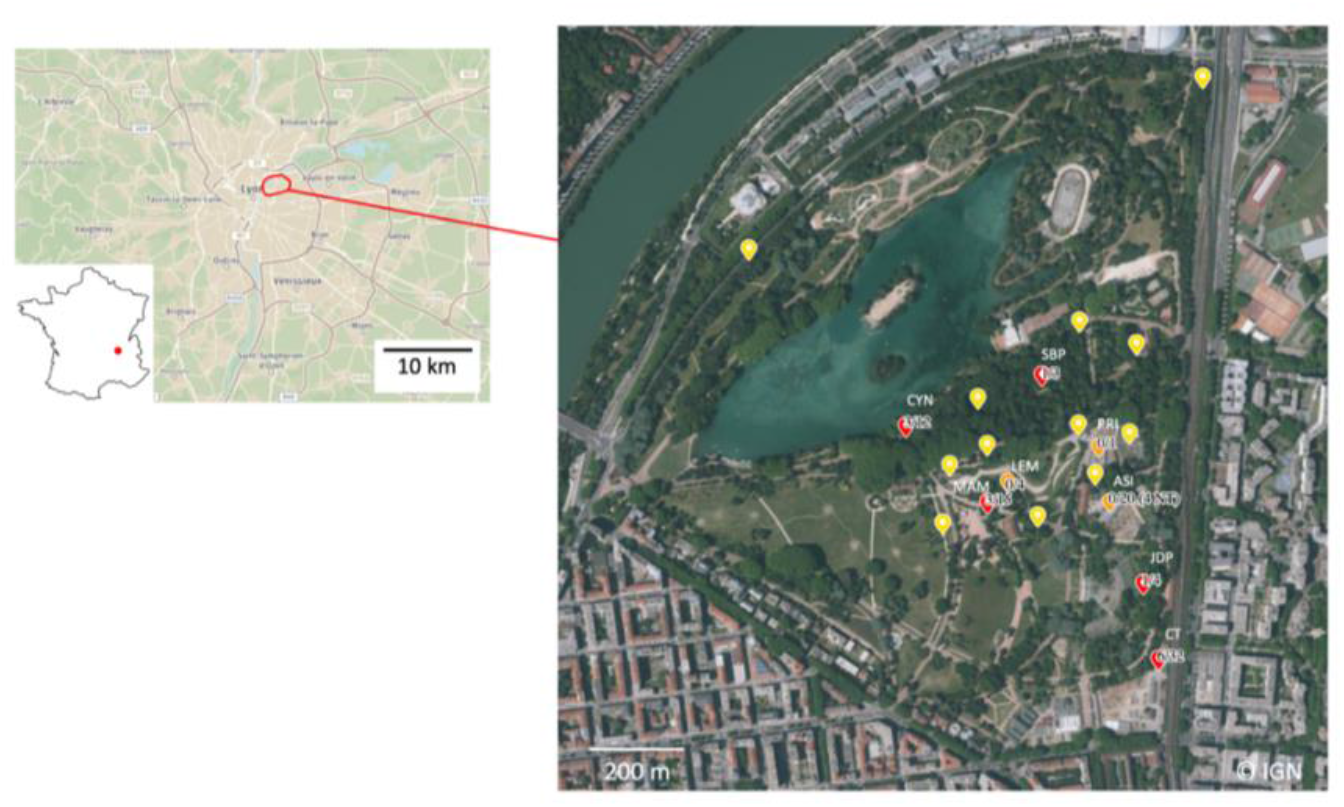
Map of the urban park of “la Tête d’Or” in Lyon, France. Yellow dots represent trap lines where no rats were caught. Orange dots represent trap lines where rats were trapped but none was seropositive for SEOV. Red dots represent trap lines where rats were trapped and some were SEOV seropositive. The codes of successful trap lines are indicated in white, and numbers indicate the proportion of SEOV seropositive rats.

### Rat population genetics

DNA was extracted from a piece of kidney stored in 96° ethanol using BioBasic kit as recommended by the supplier. Rats were genotyped with 16 microsatellite loci (D2Rat97, D3Rat159, D4Rat59, D8Rat162, D10Rat105, D11Rat11, D12Rat49, D13Rat21, D14Rat110, D15Rat64, D18Rat11, D19Rat62, D20Mit4, D1VKC1A, D1VKC1C and D5Rat43) following the procedures described in (*21*). Brown rats sampled in 2012 in Lyon city center and Givors, a city 30 km distant from Lyon, were added to the analyses.

Indices of genetic diversity and genetic differentiation were estimated using the package *hierfstat* in R v4.1.3. The population genetic structure was also analysed through a discriminant principal component analysis (DAPC) (*22*) using the package *adegenet*.

### Indirect immunofluorescence assay

The indirect immunofluorescence assay (IFA) was used to detect antibodies to orthohantavirus in 87 rat blood samples as described previously (*23, 24*). Fourteen lung samples were directed to the molecular detection.

### Library preparation and sequencing

The lung samples were homogenized, and the RNA was extracted using Trizol. Ribosomal RNA was removed using NEBNext rRNA Depletion Kit (Human/Mouse/Rat), and the sequencing libraries were prepared using NEBNext Ultra II RNA library prep kit according to the manufacturer’s instructions. The samples were sequenced using NovaSeq 6000 System (Illumina). The virus sequences were detected and annotated using Lazypipe pipeline (*25*) using fastp for trimming adapters and low -quality bases, MEGAHIT v.1.2.8 for de novo assembly and MetaGeneAnnotator for the detection of gene -like regions. These were translated to amino acids with BioPerl and queried against the UniProtKB database using SANSparallel. This was followed by re -assembly against the de-novo assembled consensus sequences using BWA-MEM algorithm implemented in HaVoC pipeline (*26*).

### Phylogenetic analyses

Phylogenetic analyses were performed on three datasets composed of complete (or semi-complete) coding regions of S, M and L segment sequences of SEOV recovered in this study and from sequences from different geographical areas available in Genbank. Sequences of Hantaan, Dobrava and Anjozorobe hantaviruses were used as outgroups in all analyses. Multiple sequence alignments were generated with the *Clustal* Omega alignment program implemented in *SeaView* v4.6.1. Phylogenetic analyses were then performed using t he Maximum Likelihood method implemented in *PhyML* 3.0 (available at http://www.atgc-montpellier.fr/phyml/) with a statistical approximate likelihood ratio test (aLRT) for branch support. The optimal substitution models were identified as the GTR+R model fo r all datasets using the “Automatic model selection by SMS” option implemented in *PhyML*. Phylogenetic trees were edited with *FigTree* v1.4.3.

Sequence identities were calculated with Sequence Identities and Similarities (SIAS) program available at (http://imed.med.ucm.es/Tools/sias.html).

### Epidemiological signature pattern analysis

The *Vespa* software (*27*) was used to detect sequence variations (amino-acids) that would be representative of the SEOV strains obtained in this study relative to a background set composed of the sequence previously characterized from a rat trapped in Lyon city (SEOV_LYON/Rn/FRA/2013/LYO852).

## Results

### Genetic analyses of captured brown rats

We successfully captured and genotyped 87 *R. norvegicus* from “la Tête d’Or”, and 107 from Lyon center and Givors, at the 16 microsatellites. Genetic diversity indices are reported in Figure 2.

**Figure 2.**
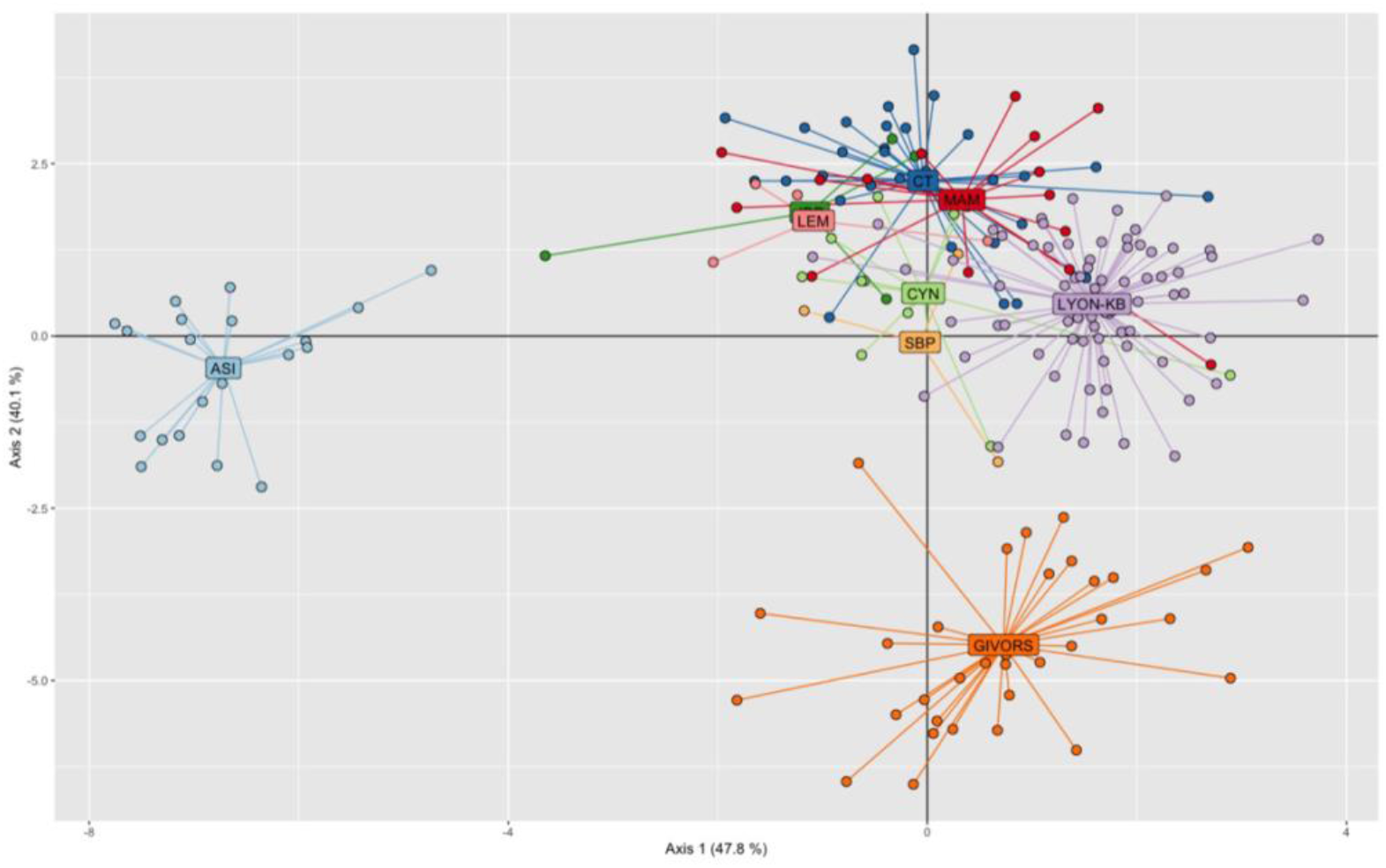
Assessment of the genetic structuring of *R. norvegicus* collected within the urban park “la Tête d’Or” in Lyon between 2020 and 2022 (trapping lines: ASI, LEM, MAM, CYN, SBP, CT, JDP, PRI), in Lyon city center (Lyon -KB) and in Givors (a city 30 km distant from Lyon), using DAPC.

Pairwise estimates of genetic differentiation (*F*_ST_) ranged from 0.00 (between MAM and SBP trapping lines) to 0.13 (between Lyon center and CT line), but they reached higher values when comparing Givors to other sampling sites (0.10 to 0.29) and more surprisingly between ASI (a trapping line within the park) and all other sampling sites (0.24 to 0.32). The DAPC confirmed the high structuring between Lyon and Givors, and the low structuring within Lyon city, between trapping lines located within (except ASI) and outside the urban park. It also showed the strong genetic differentiation between the ASI trapping line and all other sampling sites from Lyon (Figure 2).

### Detection of orthohantavirus antibodies in brown rats

The circulation of SEOV was confirmed in the French rats from the urban park in Lyon as 15 out of 87 rats were seropositive for antibodies to orthohantavirus (Table). The overall seroprevalence over the period 2020 -2022 was 17.2%. Only one rat among the 15 seropositive ones was a juvenile, so that SEOV serology was significantly influenced by rat maturity (Fisher’s exact test, *p* = 0.03). The difference in SEOV seroprevalence between males and females was not significant (X ^2^ = 1.75e-31, *p* = 1).

**Table:**
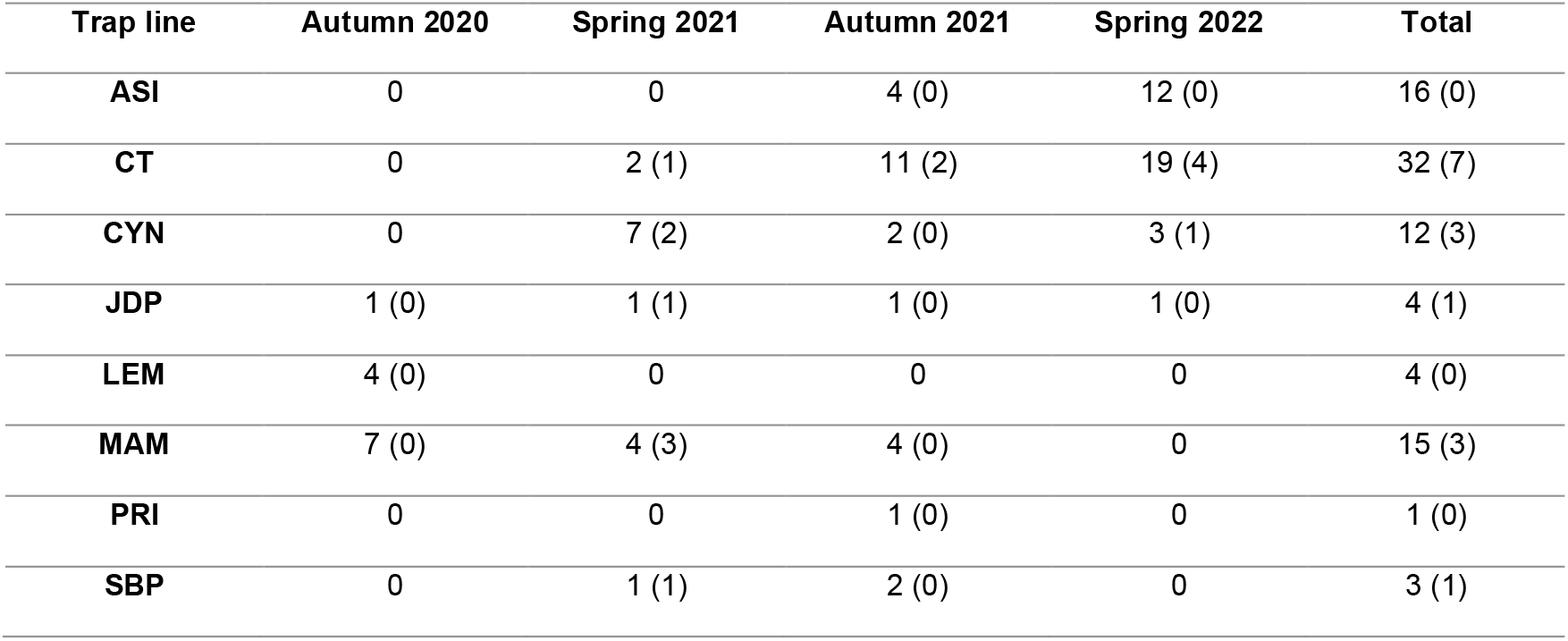
Rat captures for each trap line and trapping period. Numbers between brackets correspond to the number of seropositive rats

### Phylogenetic analysis of SEOV strains

Only the SEOV seropositive rats were sequenced on the Illumina NovaSeq 6000 Sequencing platform, which resulted in 3.1 -6.0 Gb per sample (Appendix 1). We recovered seven complete or nearly complete genomes of SEOV from those individuals, with 6453 nt for the L-segment, 3399 nt for the M-segment (all individuals), and 1287 nt for the S -segment (seven individuals).

The phylogenetic analysis showed that SEOV harbored by wild rats (*Rattus norvegicus*) in the urban park of “la Tête d’Or” in Lyon (sequences hereafter named SEOV-X-Lyon) were closely related to those of rats -derived viral genomes identified previously in Lyon city (Figure 3). All the SEOV strains from this study grouped within the previously defined lineage 7 (*13*) which includes strains from Europe (France, Belgium), South-East Asia (Singapore and Vietnam) and West Africa (Benin). The phylogenies of S, M, and L segments of the seven SEOV strains obtained in this study showed high phylogenetic proximity with strains previously collected in Lyon city (SEOV_LYON/Rn/FRA/2013/LYO852) or from its vicinity (REPLONGES|Hu|FRA|2012|12_0882). Particularly, the strain SEOV -1 (marked in red color) was closer to the sequence SEOV_LYON/Rn/FRA/2013/LYO852 than to the other strains identified in this study, whatever the phylogeny considered (S, M, or L segments, Figure 3). Interestingly, the viral genomes derived from humans and rats obtained in France clustered with lineages 7 and 9 (marked in blue), what indicates the circulation of at least two lineages of SEOV in France so far.

**Figure 3:**
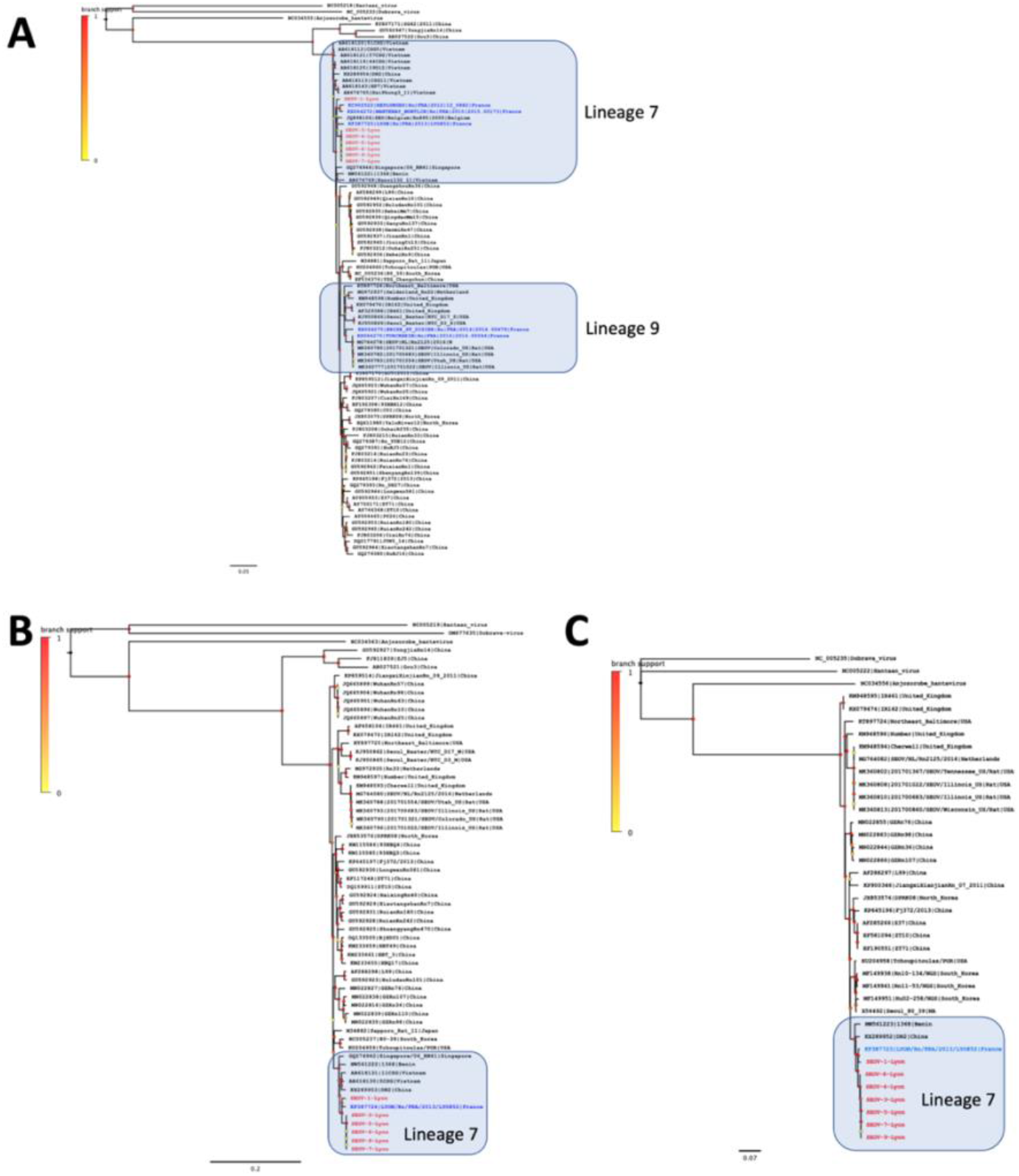
Phylogenetic analysis of SEOV gene segmen ts recovered from brown rats (*Rattus norvegicus*) trapped in Lyon city, France (in red) and reference sequences (other sequences from France are represented in blue). Phylogenetic trees were generated with PhyML 3.0 by the maximum-likelihood method on the complete coding region of the small (A), medium (B) and large (C) segments with a GTR + R nucleotide substitution model. Branch support as determined by an aLRT test is represented by a color dot at each node. Scale bars indicate n of substitutions per nucl eotide. Only lineages 7 and 9 were highlighted in the phylogenetic trees.

The SEOV strains from Lyon showed very high similarity at the amino -acid level (Figure 4). The identities were 100% for the nucleoprotein, between 99.4% and 100% for glycoprotein precursors and between 97.2 % and 100% for the RdRp. The SEOV strains detected in this study also showed high genetic similarity with the strain previously sequenced from a rat in Lyon (LYON/Rn/FRA/2013/LYO852; Identity of 100% for the nucleoprotein, 99.4% to 99.5% for the glycoprotein precursors and 98.3% to 99.8% fo r the RdRP).

**Figure 4:**
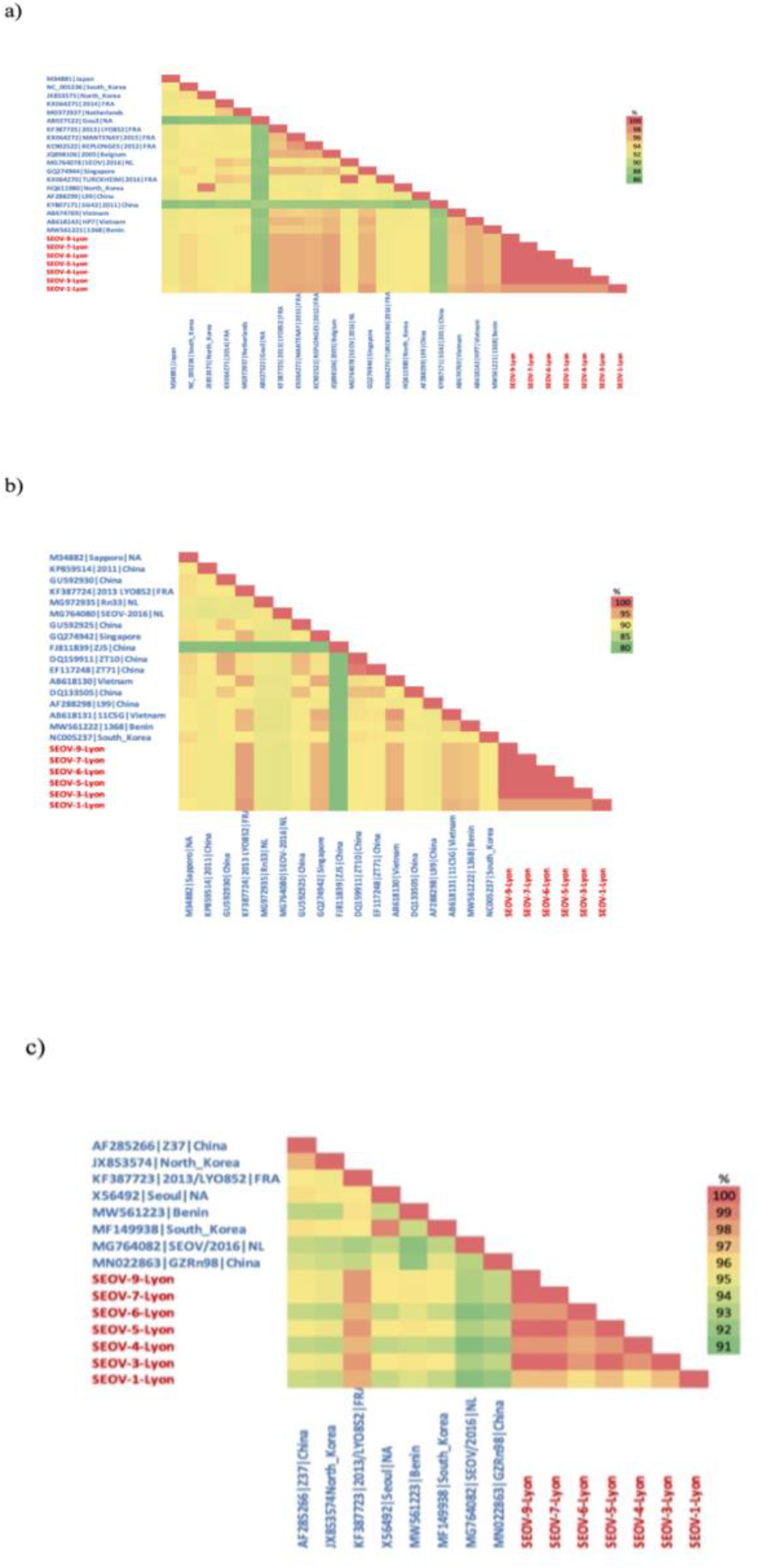
Pairwise protein sequence identities calculated with the SIAS software for a) the nucleoprotein, b) the glycoprotein precursors and c) the Rdrp. Analyses were performed considering mainly sequences belonging to lineage 7 and two SEOV -human reference sequences from France belonging to lineage 9.

### Signature pattern analysis

The signature pattern analysis enabled to identify amino acid variations that were representative of the SEOV strains detected in this study compared to the strain identified previously (SEOV_LYON/Rn/FRA/2013/LYO852). We looked for evidence of genetic variations (amino acid substitutions) of SEOV strains in Lyon between 2011 and 2021 (Appendix 2). Only mutations that were detected in all seven newly sequenced strains were considered suggestive of fixed mutations in the currently circulating SEOV strains. We did not detect any signature pattern of mutation for the nucleoprotein (see Appendix 2). For the glycoprotein precursor and RdRp, three and two mutations, respectively, were detected in all the newly sequenced strains compared to SEOV_LYON/Rn/FRA/2013/LYO852 strain (glycoproteins: V953, S1051 and L1113; RdRp: A330 and V1775).

## Discussion

Although SEOV is generally associated with a mild form of hemorrhagic fever with renal syndrome (HFRS), it still causes severe infection in some cases with a mortality rate of approximately 1-2% (*1-3*, Centers for Disease Control and Prevention). Therefore, SEOV represents a potential threat to public health. Nevertheless, SEOV is considered as an underdiagnosed virus in Europe due to the limited diagnostic assays and, presumably, unawareness among physicians. Continuous surveillance is needed to trace the virus spread, circulation and evolution in wild and domestic rats to reduce the risk of SEOV outbreak in humans.

In this context, we studied the circulation and evolution of SEOV among wild brown rats in a popular urban park in Lyon city (France), where SEOV circulation was detected ten years ago (*13*). This park is a multifacility area with a lake, a zoo, a botanical garden, restaurants, horses’ paddocks, garbage dump and carousels. Opportunities for SEOV transmission from rats to domestic animals or humans are thus numerous. It is therefore critical to pursue the eco-evolutionary surveillance of this virus, to provide an efficient early warning system and prepare risk-based interventions that target people and areas at highest risk of exposure to infected rats.

In total, 87 brown rats were caught from different sites within the urban park between 2020 and 2022. The detected overall seroprevalence (17.2%) was quite high and comparable to what has been observed in other cities (*28*). All seropositive rats except one were sexually mature, and they were trapped in several sampling sites within the park. This confirms the continuous and high level of SEOV circulation in rats in this urban park since at least 2011 (*13*), although no human case has been detected yet.

The genetic analyses of rats have revealed a main cluster consisting of rats from both the park and the city center of Lyon. This suggests a large rat population and possibility of gene flow between the park and the rest of the city. As a result, SEOV may potentially circulate and evolve throughout Lyon, so that SEOV surveillance should be conducted at the scale of the city.

Interestingly, rats trapped in the ‘Asian forests’ building of the zoological park (ASI), an area opened in 2021 to home a large diversity of mammals and birds, showed high levels of genetic differentiation compared to the rats trapped in other parts of the park. They were all seronegative. This could suggest a recent introduction, or a strong founder effect associated with the creation of this area. It should be important in the future to pay particular attention to this site, to survey the evolution of rat populations and SEOV prevalence in rats and captive monkeys (*Nomascus leucogenys*) of the ‘Asian forests’ building.

We recovered seven complete and semi-complete genomes of SEOV from these wild rats, which greatly increase the genetic data available on SEOV from France, especially for the M and L segments. The recovered genomes were significantly larger in contrast to the previous studies (*11-13*). Genetic analyses showed high identity levels between all these sequences. Indeed, only a few amino acid substitutions were identified between 2011 and 2021, on the glycoprotein precursor and the RdRp. This suggests strong purifying selection as already described (*29*). Note that the lack of complete genomes available prior to this study requires considering these results with caution.

Phylogenetic analyses revealed that all SEOV strains retrieved in this study clustered together with the recent SEOV strain from Lyon (SEOV_LYON/Rn/FRA/2013/LYO852) with SEOV-1 being slightly more distant from the others. This shed the light on the origin of SEOV strains circulating in France and in Lyon city. The strains described in this study clustered together with strains from Asia (Vietnam, Singapore, and China) and the recently sequenced one from Africa (Benin) within the SEOV lineage 7 (*21*). South-East Asia was suggested to be the origin of lineages 7 and 9 that are circulating in Europe and causing infections in humans (*30*). Both lineages were identified in France (*11*), with a human case associated with lineage 7 near Lyon (*12*), suggesting multiple SEOV introduction events in France. In the present study, we observed only strains from lineage 7 circulating in Lyon.

Taken together, these findings suggest that the same strain of SEOV has been circulating continuously in Lyon since at least 2011 at a significant seroprevalence, with only a few new fixed mutations detected in 2020 -2022. Our results underscore the importance of regularly monitoring SEOV strains present in rats in urban areas like Lyon. Enhanced surveillance of SEOV transmission among rats can assist in better preventing and identifying potential outbreaks in humans, which is critical in mitigating the associated zoonotic risk.

## Acknowledgements

We thank all the people from the Health department, the park “la Tête d’Or’ and the zoological park of Lyon city (especially Cédric Verjat, Alban Chauvet and Fabrice Delaveau) for their help in the field.

## Biographical Sketch

Alburkat Hussein is a virologist and PhD researcher at the University of Helsinki, faculty of medicine, Helsinki, Finland. His research interest includes next generation sequencing, hantaviruses, arenaviruses, and virus-host interactions.

## Appendixes

**Appendix 1:** Results provided by the NovaSeq platform.

**Appendix 2:** Frequencies of amino-acids of the query set sequences (SEOV strains derived from the present study) and the background sequences (SEOV_LYON/Rn/FRA/2013/LYO852 sequences derived from the database GenBank). a) Signature of amino acids detected in the SEOV M -segment. b) Signature of amino-acids detected in the SEOV L-segment. The first upper line shows the query signature amino -acids and the two lines below show the frequency of those amino -acids among the query set and the background set, respectively. The fourth line illustrates the common amino-acids detected among the background set. The following two lines beneath show the frequency of those amino-acids among the query set and the background set sequences, respectively. The last line refers to the alignment position among those sequences. The figures under each table show the amino-acid variations

