## Appendix 1 for "Evolution and Genetic Characterization of Seoul Virus in Wild Rats *Rattus Norvegicus* from an urban park in Lyon, France 2020-2022"

**Appendix 1:** Results provided by the NovaSeq platform.

|  | Total reads | Reads passing filter | Mapped to L | Mapped to M | Mapped to S |
| --- | --- | --- | --- | --- | --- |
| SEOV-1 | 21647992 | 20616344 | 1922 | 3368 | 900 |
| SEOV-2 | 19955700 | 18572006 | 0 | 0 | 0 |
| SEOV-3 | 17070824 | 16346840 | 6264 | 14376 | 4400 |
| SEOV-4 | 15143328 | 14378442 | 2360 | 4860 | 1670 |
| SEOV-5 | 13791536 | 13116448 | 4494 | 8568 | 2366 |
| SEOV-6 | 13663814 | 12977102 | 1072 | 2500 | 1894 |
| SEOV-7 | 16468838 | 15785140 | 4544 | 8608 | 4596 |
| SEOV-8 | 19754154 | 18802908 | 0 | 0 | 0 |
| SEOV-9 | 23957098 | 22871504 | 9587 | 15812 | 8172 |
