## Supplementary figures and images for "Evolution and Genetic Characterization of Seoul Virus in Wild Rats *Rattus Norvegicus* from an urban park in Lyon, France 2020-2022"

### Appendix 2A

#### Slide 1
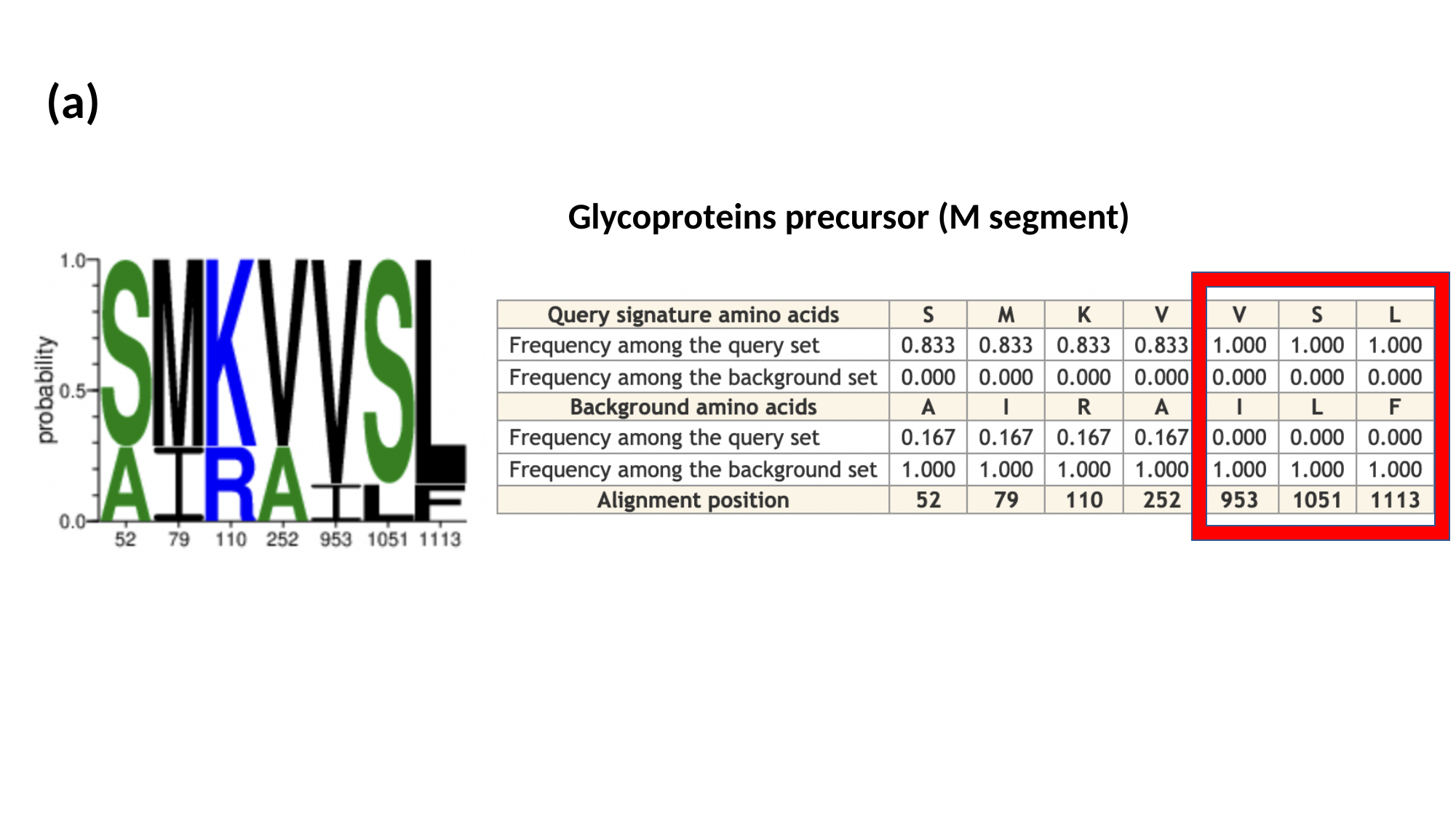

(a)
Glycoproteins precursor (M segment)

### Appendix 2B

#### Slide 1
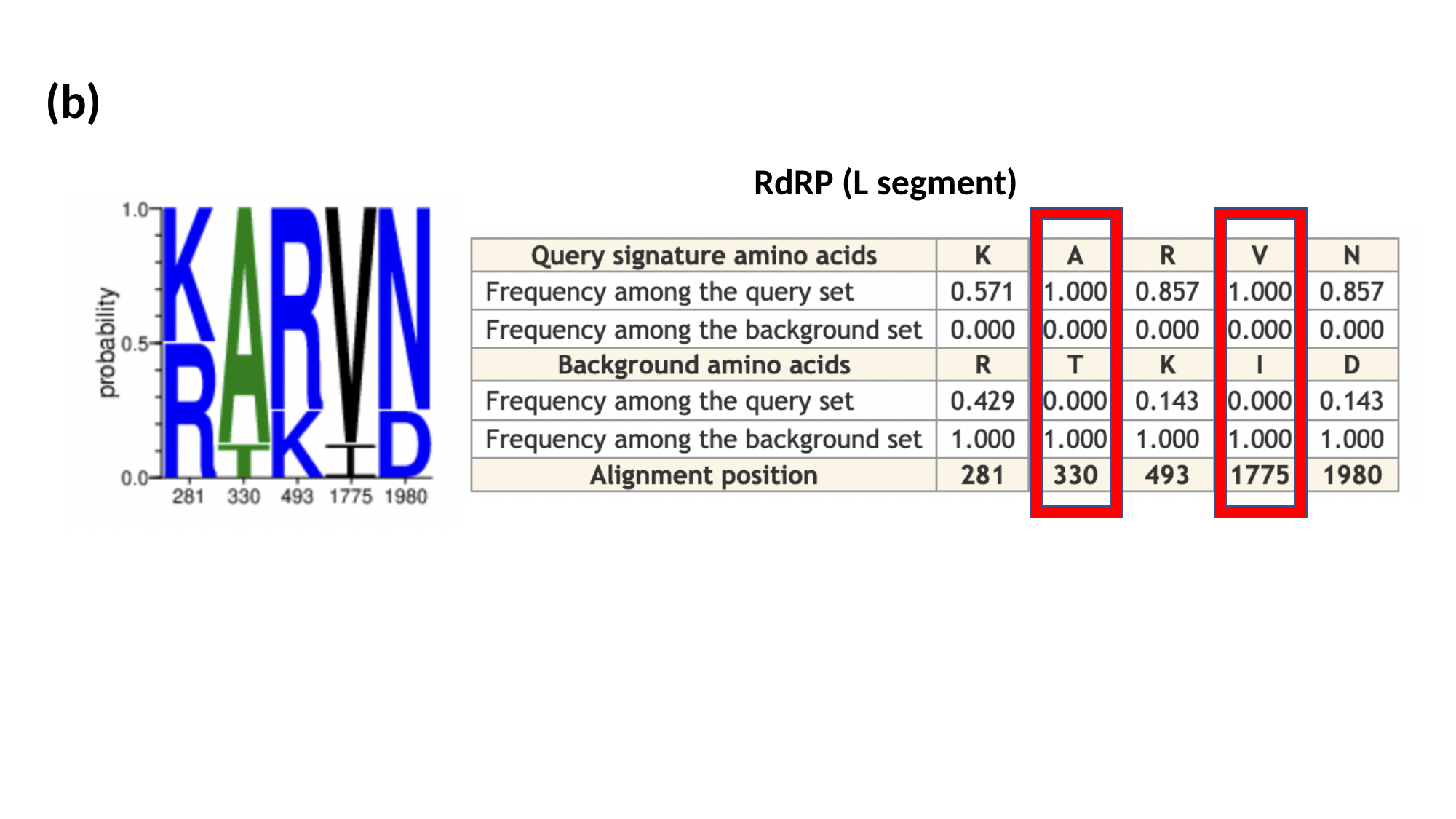

(b)
RdRP (L segment)
